# Near-infrared Spectroscopy evaluations for rapid differentiation of carbapenem resistant Enterobacteriaceae from susceptible strains

**DOI:** 10.1101/2020.08.06.240804

**Authors:** Bushra Alharbi, Maggy T. Sikulu-Lord, Anton Lord, Hosam M Zowawi, Ella Trembizki

## Abstract

Antimicrobial resistance (AMR) caused by Carbapenem-Resistant Enterobacteriaceae (CRE) is a global threat. Accurate identification of these bacterial species with associated AMR is critical for their management. While highly accurate methods to detect CRE are available, they are costly, timely and require expert skills making their application infeasible in low-resource settings. Here, we investigated the potential of Near-infrared Spectroscopy (NIRS) for a range of applications; i) the detection and differentiation of isolates of two pathogenic Enterobacteriaceae species, *Klebsiella pneumoniae* and *Escherichia coli* and, ii) the differentiation of carbapenem resistant and susceptible *K. pneumoniae*. NIRS has successfully differentiated between *K. pneumoniae* and *E. coli* isolates with a predictive accuracy of 89.04% (95% CI; 88.7-89.4%). *K. pneumoniae* isolates harbouring carbapenem resistance determinants were differentiated from susceptible *K. pneumoniae* strains with an accuracy of 85% (95% CI; 84.2-86.1%). To our knowledge, this is the largest demonstration of a proof of concept for the utility and feasibility of NIRS for rapidly differentiating between *K. pneumoniae* from *E*.*coli* as well as from carbapenem resistant *K. pneumoniae* from susceptible strains.

## Introduction

Infections caused by Carbapenem-Resistant Enterobacteriaceae (CRE) are emerging as a global health concern. They are associated with difficulties in treatment and a major contributing factor to global morbidity and mortality (1). Carbapenem-resistant pathogens are also listed as a critical priority in the World Health Organization global Priority Pathogens List (1), which primarily include *K. pneumoniae* and *E*.*coli*. Performing accurate, efficient and fast detection of CRE in clinical laboratories is a key factor to antimicrobial stewardship and appropriate management of patients. Access to affordable and high throughput diagnostics for surveillance of CREs is also needed, particularly in low resource settings (2). Various techniques are currently used in routine clinical diagnosis and surveillance to identify species and ascertain AMR. These may depend on the settings and include traditional phenotypic methods such as biochemical tests and gold standard bacterial culture methods (3).

Additionally, molecular methods, including commercial PCR-based platforms and Whole Genome Sequencing (WGS), which have revolutionised clinical diagnostics and play a significant role in bacterial typing (4). However, these methods are costly, time and labour intensive. Consequently, the practical application of these methods is not feasible in resource limited settings where disease burden is high and where syndromic-based diagnosis is the mainstay (5-10). Ultimately, a simple, cost-effective, rapid and reproducible alternative for easy identification and characterisation of bacterial isolates and/or clinical samples should be applied.

Near Infrared Spectroscopy (NIRS) is a technique that uses the near-infrared region of the electromagnetic spectrum (700–2500nm) to characterise biological samples based on a reflected spectral signature. The spectral signature is collected following the interaction of biological samples with infrared light (11). The resultant spectral signature is unique for various biological samples based on the type of their chemical profile. NIRS is rapid and non-invasive as well as a simple technique requiring little to no sample preparation procedures and or reagents to operate (11). NIRS is applied in multiple fields such as agriculture (e.g. assess food quality and safety, and detection of seeds germination and viability) (12-14), food microbiology (e.g. assess contamination) (15), medical research (e.g. non-invasive diagnosis and pathophysiology) (16), entomology (e.g. to detect viruses in mosquitos) (17-19) and chemistry (e.g. measuring chemical properties of matters) (20).

There are only a handful of studies exploring NIRS to differentiate resistant from susceptible strains and one species from another (21-25). The data so far is encouraging yet limited by sample size or insufficiently characterised sample banks using well-established reference methods. In addition, the variability in data analysis approaches (i.e. machine learning algorithms) and sample preparation make it challenging to compare and assess further and reproducible utility. Accordingly, in this study we aim to further close the gap and elucidate NIRS feasibility in this arena. Here, we applied NIRS on a unique well-characterized *K. pneumoniae* and *E. coli* sample banks from countries in the Middle East to i) differentiate *K. pneumoniae* from *E*.*coli* and ii) evaluate its ability to differentiate between wild type *K. pneumoniae* from carbapenemases producing strains.

## Materials and Methods

### Bacterial Isolates and sample preparation

Two bacterial species were used for this experiment and are described in detail in **Table 1**. ; *E*.*coli* (N=40) and *K. pneumoniae* (N=73). Clinical isolates were originally collected from Saudi Arabia (*E*.*coli* n= 2; *K. pneumoniae* n=40), Bahrain (*K. pneumoniae* n=1), Qatar (*E*.*coli* n= 4; *K. pneumoniae* n=5), Oman (*E*.*coli* n= 2; *K. pneumoniae* n=3), United Arab Emirates (*E*.*coli* n= 8; *K. pneumoniae* n=6), Jordan (*E*.*coli* n=19; *K. pneumoniae* n=10), Egypt (*E*.*coli* n= 5; *K. pneumoniae* n=7), Syria (*K. pneumoniae* n=1). Bacterial species were confirmed by matrix-assisted laser desorption ionization–time of flight mass spectrometry (MALDI-TOF MS) on a Microflex platform (Bruker Daltonics, Inc.) (26). Initial screening for carbapenem resistance was determined by measuring reduced susceptibility to ertapenem by disk diffusion; and DNA extracts of the isolates were PCR tested for the presence of carbapenemases determinants (*bla*_NDM_, *bla*_OXA-48_. *bla*_KPC_, *bla*_VIM_, and *bla*_IMP_ types) (27).

**Table 1.**
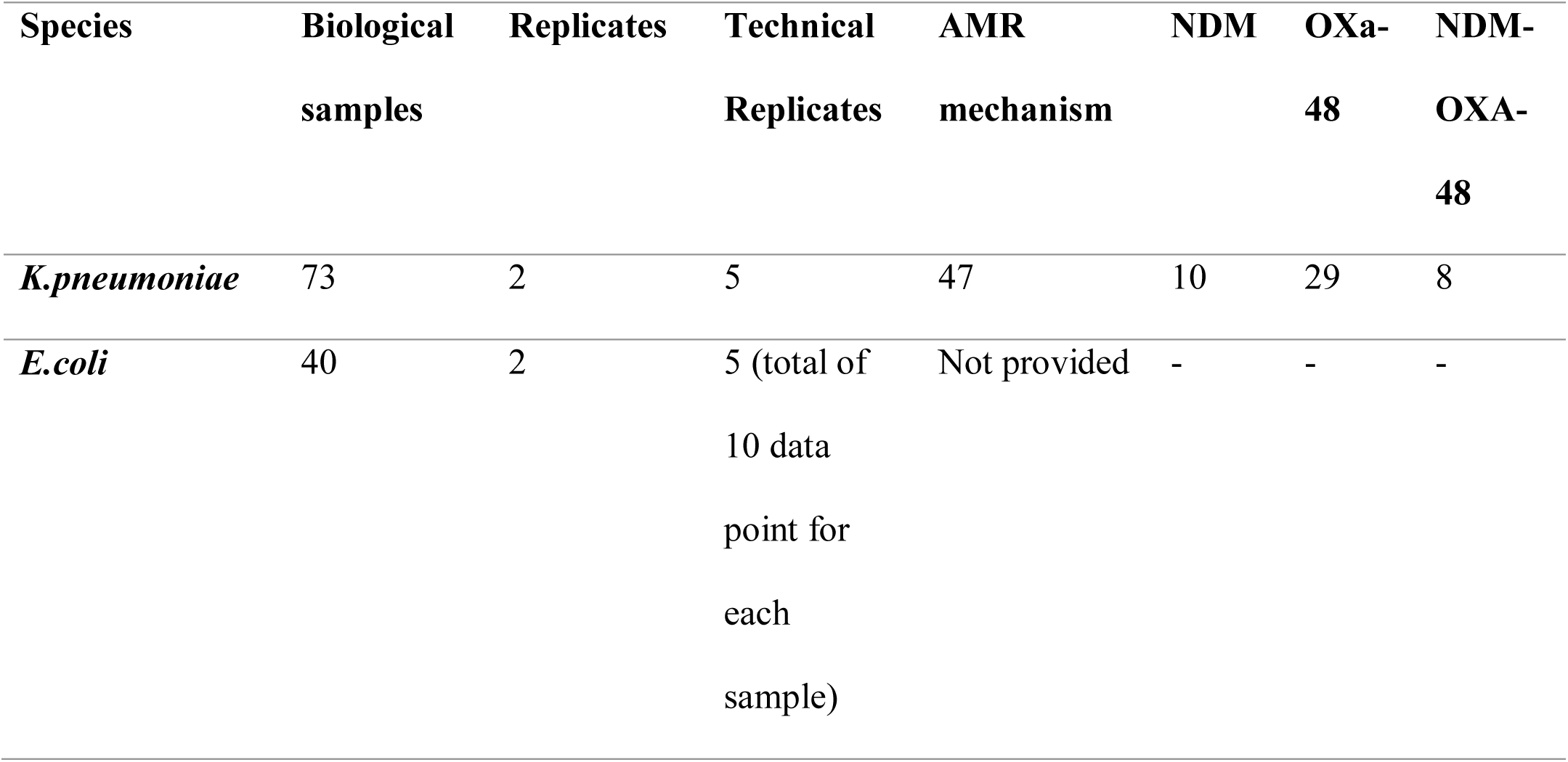
Summary of samples describing the resistance mechanisms of the bacteria and the number of replicates including biological and technical replicates.

| Species | Biological<br>samples | Replicates | Technical<br>Replicates | AMR<br>mechanism | NDM | OXA-48 | NDM-OXA-48 |
| --- | --- | --- | --- | --- | --- | --- | --- |
| *K.pneumoniae* | 73 | 2 | 5 | 47 | 10 | 29 | 8 |
| *E.coli* | 40 | 2 | 5 (total of<br>10 data<br>point for<br>each<br>sample) | Not provided | - | - | - |

### Molecular confirmatory analysis of bacterial resistance determinants

All *K. pneumoniae* samples (n=73) were further tested by a multiplex PCR method previously described (SpeeDx Pty Ltd., Australia) (27). Briefly, samples were screened for carbapenemase genes *bla*_KPC_, *bla*_NDM_, *bla*_OXA-48-like_, *bla*_IMP-4-like_ and *bla*_VIM_ in a single multiplex reaction. Reactions were amplified using ABI7500 real-time PCR instrument (ThermoFisher Scientific, Australia) with cycling conditions; an initial 95 °C 2 min hold, followed by 10 touch-down cycles at 95 °C for 5 s and 61 °C (− 0.5 °C per cycle) for 30 s, followed by 40 cycles at 95 °C for 5 s and 52 °C for 40 s.

### Scanning with NIRS

All isolates were sub-cultured twice on Mueller Hinton (MH; Becton Dickinson and company, France) plates and incubated for 12 hours at 37 °C before processing for NIRS analysis. Following 24 hours incubation, bacterial isolates were inoculated into 2 ml of deionised water at a cell density of 4.0 McFarland. Approximately between 5 technical replicates of 3 μL of each bacterial suspension were placed on microscopic glass slide and were left to dry for approximately 10 minutes before scanning with the NIRS instrument. The dried spots were scanned by Labspec 4*i* NIR spectrometer (Malvern Panalytical, Malvern, United Kingdom) with wavelengths ranging from 350 to 2350 nm using a fiber optic probe with 6 illumination fibres. 40 biological samples of *E. coli* (with 2 replicates each; n=80) and 73 biological samples of *K. pneumoniae* (with 2 replicates each; n=146) scanned by NIRS. These were further split into 5 technical replicates for each sample, resulting in 10 data points for each biological sample (Table1.).

### Data pre-processing

The absorbance spectral data generated from the labspec 4i were converted to reflectance spectra using equation 1. Each spectra was mean centred and normalised for variance (28, 29). Baseline correction and variance normalisation were performed using the caret package in R 3.5.1. Outcomes (e.g. species, resistance) were coded in a binary format (0 or 1) for each classifier and predictions were generated on a continuous scale. Predictions above 0.5 were considered to belong to the class ‘1’, while predictions below 0.5 were considered to belong to the class ‘0’. The individual and average spectra have undergone pre-processing steps, including mean centering and variance normalisation to decrease the noise and improve signal-to-noise ratio in the spectra.

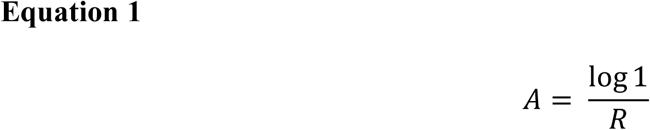

Where A = absorbance and R = reflectance

### Model development and calibration

Predictive models were developed using the partial least squares (PLS) method using the ‘pls’ package implemented in ‘R’ software (3.5.1) (30). K-fold cross validation was used (k=10) to validate the model. That is, data was divided into 10 groups, for each run, 9 sets of data were used to train the model with the last group used to test the accuracy. The maximum number of regression factors for each model was 20. The number of factors used in each model was chosen based on the lowest number of factors required to reach the maximum accuracy within the training dataset. This process was repeated 10 times until each group had been held out once. Reported statistics are for the testing groups only. Three classification models were developed to differentiate; (1) *E. coli* from *K. pneumoniae*, and (2) *K. pneumoniae* carbapenem resistance-gene-positive from *K. pneumoniae* carbapenem resistant-gene-negative. Each of the models was then applied to predict the identity of samples that were not used in training the model. Accuracy, sensitivity and specificity were calculated by comparing the results against the reference methods for bacterial species confirmation and carbapenem genes detection.

## Results

### Differentiation of *E. coli* and *K. pneumoniae*

Using PLS *E*.*coli* and *K. pneumoniae* were differentiated with an accuracy of 89.05% (95%CI 88.7-89.4%, < 0.0001) (N = 113). Sensitivity and specificity for differentiating the two species were 92.7% and 84.7%, respectively (**Table 2**). To ensure the validity of these results K-fold cross validation (k=10) was used. Results presented here are for the testing set.

**Table 2.** Results summary for accuracy, specificity and sensitivity for each of study analysis

| Classification<br>Model | Accuracy% | Specificity<br>% | Sensitivity% | p-value | Total sample<br>Numbers (N) |
| --- | --- | --- | --- | --- | --- |
| *E.coli* and *K.*<br>*pneumoniae*<br>differentiation | 89.04% | 84.74% | 92.75% | <0.0001 | 114 |
| *K. pneumoniae*<br>resistant vs.<br>wild type<br>differentiation | 85% | 81% | 89% | <0.0001 | 73 |

Figure 1-A illustrates the normalised spectra in the region of 700-2350 nm for *E*.*coli* and *K* .*pneumoniae*. Accordingly, a PLS with cross-validation diagnostic method was used to develop the calibration model. In the PLS model, the analysis was conducted as binary format (1, 0) in a density plot (Figure 1-B). A value of “0” was assigned to *K. pneumoniae* and a value of “1” was assigned to *E. coli*. Overlapping between the two data (Pink colour was used to represent *E. coli* and Blue to represent *K. pneumoniae*) of the continuous interval were considered as misclassified.

**Fig. 1.**
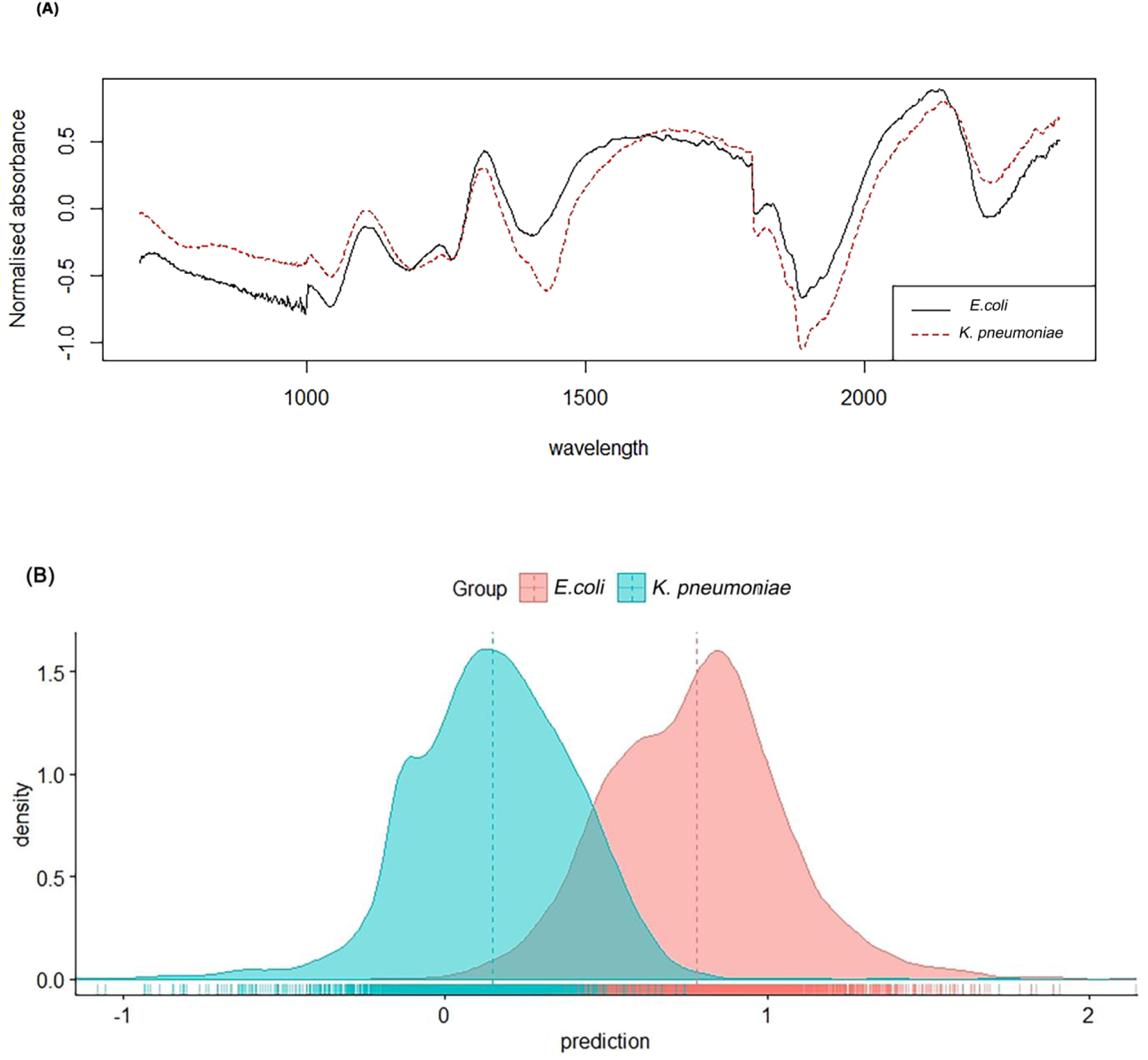
**(A)** Average NIR spectra in the 350 to 2500 nm region from *K. pneumoniae* (red) and *E. coli* (black) and **(B)** Density plot showing NIRS differentiation of *E*.*coli* (pink) and *K. pneumoniae* (Blue) using test samples.

### Differentiation of susceptible and resistant *K. pneumoniae* using NIRS

Further analysis was conducted to predict susceptibility and resistance among *K. pneumoniae* samples. The samples were previously characterised and classified as susceptible/not detected AMR mechanisms (n=29) or resistant [with OXA-48 (n=28), NDM (n=10)], and OXA-48 with NDM (n=6)]. In this analysis, we evaluated NIRS for its ability to differentiate susceptible *K. pneumoniae* from resistant samples regardless of the mechanism of action. The PLS model resulted in an accuracy, sensitivity and specificity of 85% (95% CI; 84.16-86.06%, p < 0.0001), 89% and 81%, respectively (**Table 2**.). Figure 2-A illustrates the normalised average spectra of resistant and susceptible *K. pneumoniae*. Similarly to the above PLS model analysis, conducted as binary (1, 0) in density plot **Fig. 2 (B)** a value of “0” was assigned to resistant *K. pneumoniae* and a value of “1” was assigned to susceptible *K. pneumoniae*. Overlapping between the two data of the continuous interval were considered as misclassified (Pink colour was used to represent carbapenem susceptible, while Blue colour was used to represent carbapenem resistant strains) **Fig. 2 (B)**

**Figure 2.**
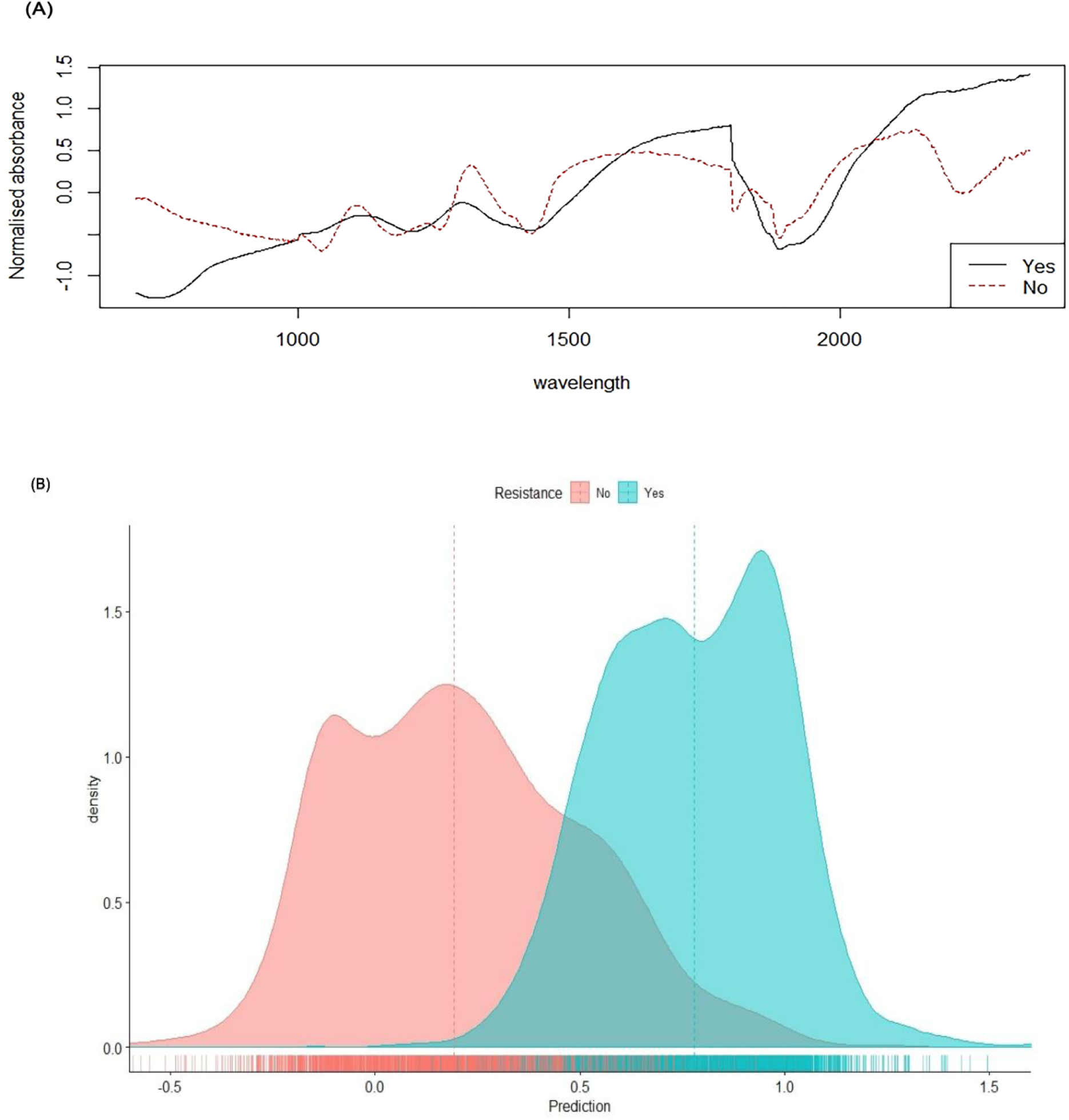
**(A)** Normalised NIR spectra in the 350-2500 nm region from susceptible *K. pneumoniae* (red line) and resistant *K. pneumoniae* (black line) Panel **(B)** Density plots showing NIRS differentiation of *K. pneumoniae* AMR-genes negative or susceptible (Pink) and AMR harbouring or resistant (Blue)

## Discussion

The overall objective of this study was to explore the applicability of NIRS to differentiate between *E*.*coli* and *K. pneumoniae*, and to differentiate between *K. pneumoniae* harbouring AMR genes from strains that absent for AMR genes (or otherwise, wild type). Here, we show that NIRS has the potential to differentiate these species with a predictive accuracy of 89% and can predict certain carbapenemase-encoding genes with an accuracy of 85%. Specificity and sensitivity for differentiating species (*E. coli* and *K. pneumoniae*) was 85% and 92%, respectively and specificity and sensitivity for the AMR-gene harbouring vs. wild type (*K* .*pnuemoniae*) strains was 81% and 89%, respectively.

Spectroscopy techniques to identify clinical bacteria is an emerging technique in the medical field but a widely applied in the food industry (31). However, only three studies have previously explored the differentiation of bacterial species utilising the NIRS technique that can be assessed against our study. Although sample preparation, sample size or machine learning techniques across these studies differed, predictive accuracies obtained are comparable to our results. One study utilised a miniature portable Fourier-Transform NIR spectrometer (900-2600 nm) to differentiate *bla*_KPC-2_-harbouring from *bla*_KPC-2_-negative *K. pneumoniae* clinical isolates by collecting spectral signatures of bacteria DNA on aluminium-plated backing plate. Algorithm-linear discriminant analysis (GA-LDA) and successive projection algorithm (SPA-LDA) models were used to analyse spectral data. Predictive sensitivity using GA-LDA and SPA-LDA for *bla*_KPC_-negative was 100% and 76%, respectively compared to the predictive sensitivity of 66% for *bla*_KPC-2_-harbouring *K. pneumoniae* using either model (25). These data are comparable to our findings where we demonstrated that sensitivity of NIRS for predicting *bla*_NDM-type_ and *bla*_OXA-48-type_-genes harbouring *K. pneumoniae* was slightly lower (81%) than that of wild-type (92%).

A plausible limitation for the differentiation of resistant and susceptible strains in our study is the potential that the organism harbours additional resistance determinants or variations which were not previously characterized resulting in a ‘false negative’ results. Alternatively, the detection of a gene which is not expressed, in-turn resulting in ‘false positive’ result. This reinforces the need for further validation and testing on large well-characterized (genotype and phenotype) sample banks to best account for such variations.

Kammies and colleagues investigated the use of NIRS hyperspectral imaging within the wave regions of 900-2500nm to detect and differentiate *Bacillus cereus* and two *Staphylococcus strains (aureus* and *epidermidis)*. Samples were streaked onto solid Luria-Broth (LB) and spectra was collected directly from the petri-dishes. Data was analysed with PLS and a predictive accuracy of 90.98% (95%CI; 82-99.96%) was achieved (22). Lastly, when NIR within the wavelengths of 750 to 1350 nm was used to differentiate two food-borne *E. coli* strains, ATCC 25922 (n= 5) and K12 (n= 5) grown in liquid media and Artificial Neural Network (ANN) was used to analyse spectral data, a regression coefficient (R^2^) of 0.98 was achieved (23).

Here we applied PLS to differentiate the two species and to detect resistance with high predictive accuracies. It is indeed possible that other machine learning techniques would generate an improved result, however the sample size used in our study was best suited for PLS analysis. We recommend that future work with a relatively larger sample size should explore the possibility to employ other machine learning techniques for data analysis.

Finally, we demonstrated for the first time that NIRS can rapidly differentiate, with reasonable accuracy, between resistant and susceptible *K. pneumoniae* strains harbouring a range of common AMR associated mutations. Further studies are required to assess NIRS feasibility for the identification and differentiation between and within bacterial species and strains. Future work would include evaluating additional machine learning algorithms, increased samples size and variably, limit of detection studies, culture media comparisons to determine the effects of noise background, and finally, evaluate and develop a protocol for screening directly from clinical samples (i.e. non-culture). Importantly, a side-by-side evaluation of NIRS with Whole Genome Sequencing and phenotypical antimicrobial susceptibility data would be most advantageous for a meaningful comparable data set.

## Conclusion

To our knowledge, this is the largest evaluation of NIRS feasibility in differentiating *K. pneumoniae* from *E. coli*, and differentiating *K. pneumoniae* carbapenem resistant from susceptible strains. This proof of concept demonstrates the potentiality of NIRS in microbial identification and AMR characterisation.

## Ethics Statement

The bacterial isolates that used for this project were stored at the UQCCR biobank. These samples were originally isolated from clinical samples shipped from Middle East countries. The bacterial samples received ethics approval; they were covered by human ethics clearance number 2018000615.

## Acknowledgments

ET holds a National Health & Medical Research Council Early Career Fellowship HMZ was kindly supported by The Marchant Foundation Fellowship. Part of this work was presented as a Poster presentation at International conference for Near Infrared Spectroscopy, NIR 2019, September 2019 Gold Coast, Australia as well as in the Gulf Cooperation Council Microbiology and Infectious Diseases Conference 2019 in Dubai, and the abstract was published in the journal of infection and public health issue (32). The NIR instrument used for this study was provided by the United States Department of Agriculture (USDA) through Dr Floyd Dowell. Mention of trade names or commercial products in this publication is solely for the purpose of providing specific information and does not imply recommendation or endorsement by the U.S. Department of Agriculture. USDA is an equal opportunity provider and employer.

**Bushra Alharbi:** Laboratory testing, Writing Original Draft **Maggy Sikulu-Lord:** Supervision, Writing - Review & Editing **Anton Lord:** Formal statistical analysis, visualization, manuscript editing **Hosam M Zowawi:** Conceptualization, sample bank, editing of manuscropt **Ella Trembizki:** Conceptualization, Resources, Supervision, Writing - Review & Editing, and Project administration.

## Conflict of interest

All authors declare no conflict of interest

All data needed to evaluate the conclusions in the paper are present in paper and/or the Supplementary Materials. Additional data related to this paper may be requested from the authors.

